# Inference of disease-associated microbial gene modules based on metagenomic and metatranscriptomic data

**DOI:** 10.1101/2021.09.13.460160

**Authors:** Zhaoqian Liu, Qi Wang, Anjun Ma, Dongjun Chung, Jing Zhao, Qin Ma, Bingqiang Liu

## Abstract

The identification of disease-associated microbial characteristics is crucial for disease diagnosis and therapy. However, the heterogeneity, high dimensionality, and large amounts of microbial data present tremendous challenges for the discovery of key microbial features. In this paper, we present IDAM, a novel computational method for disease-associated gene module inference from metagenomic and metatranscriptomic data. This method integrates gene context conservation (uber-operon) and regulatory mechanisms (gene co-expression patterns) to explore gene modules associated with specific phenotypes using a mathematical graph model, without relying on prior meta-data. We applied IDAM to publicly available datasets from inflammatory bowel disease, melanoma, type 1 diabetes mellitus, and irritable bowel syndrome and demonstrated the superior performance of IDAM in disease-associated characteristics inference compared to popular tools. We also showed high reproducibility of the inferred gene modules of IDAM using independent cohorts with inflammatory bowel disease. We believe that IDAM can be a highly advantageous method for exploring disease-associated microbial characteristics. The source code of IDAM is freely available at https://github.com/OSU-BMBL/IDAM.

## Introduction

Trillions of microbes colonize the human body and play critical roles in multiple fundamental physiological processes, and changes in microbial composition and functions are intimately interwoven with multiple diseases, ranging from obesity to cancer and autism^1,2^. With the rapid development of sequencing technologies, the Human Microbiome Project^3^, integrative Human Microbiome Project^4^, and Metagenomics of the Human Intestinal Tract project^5^ have generated large amounts of microbial data related to numerous diseases. Considerable effort has been aimed at using these data to produce clinical insights and deepen the understanding of the mechanisms responsible for disease progression and treatment^6^. Some pathogenic organisms have been found to be closely related to disease states and can be used as potential markers of disease occurrence and development^7,8^. These findings hint at a promising avenue for clinical application—disease diagnosis and therapy—via the use of key microbial taxonomic or functional characteristics.

One of the classical paradigms of key microbial characteristics identification is to detect taxa and metabolic pathways that can statistically significantly distinguish between two or more groups^9-11^. Additionally, given the success of machine learning in multiple scenarios, there have been several attempts to use machine learning in the identification of microbial characteristics^12,13^. Such methods include random forest and deep feedforward networks^12,13^. However, these existing studies face several challenges. *First*, most of them focus on 16S ribosome RNA and metagenomic (MG) data. 16S ribosome RNA data suffers from low taxonomic resolution and an absence of functional information^14^, which may induce spurious characteristic identification with diseases since diverse microbial communities from different patients can have very similar functional capabilities^14^. While MG data can provide information at all taxonomic levels, researchers have found that a large proportion of reads cannot be successfully mapped to existing reference genomes during taxonomic identification^15,16^. Furthermore, only a minority of genes are well-annotated with biochemical functions^15^. The limitations of reference genome databases and annotation systems will lead to the potential loss of valuable information while taxonomic or metabolic pathways analysis using MG data^15,16^. Meanwhile, the functional analyses produced by MG analyses are not necessarily equivalent to the true activities within the microbiome. Multiple metagenomically abundant genes in the human gut microbiome are significantly down-regulated at the transcriptional level^17^. *Second*, existing methods are designed based on the strong assumption that data with sufficiently large sample sizes and accurate metadata information are available to design groups or train models. However, the current metadata related to many sequencing samples is incomplete, misleading, or not publicly available^18^, which may lead to these methods being infeasible or causing bias in key characteristic inference. These challenges drive the need to develop an easy-to-use and effective method that keeps up with current data for disease-associated microbial characteristic inference.

In this study, we proposed a novel method to **I**nfer **D**isease-**A**ssociated **M**icrobial gene modules (**IDAM**), which to a large extent overcomes these limitations. The data we used was matched MG and metatranscriptomic (MT) data sequenced from microbial samples for assessing active functions within communities^19^. To make full use of the available data, we focused on a community-level analysis that goes beyond species level^15^, so that the reads do not need to be mapped to reference genomes, and provided a comprehensive overview of functions performed by either individual or different microbial assemblages.

We investigated disease-associated key characteristics by detecting gene modules with specific co-expression patterns across certain subsets of samples^19^, without requiring prior metadata of samples. This can be a complement to existing supervised methods in the absence of high-quality metadata. Here, we noted that one cannot fully characterize complex functional activities based solely on ensemble co-expression, by which gene expression is assessed using reads from various species (**Fig. 1a**) and genes with no biological relevance may be clustered together (**Fig. 1b**). Fortunately, previous studies have suggested that functionally associated genes are generally highly conserved during evolution^20,21^, forming uber-operon structures (**Supplementary Fig. S1**). This finding motivated us to integrate gene context conservation to study cross-species functional-related mechanisms, thereby alleviating the bias in key microbial characteristics inference. This idea has not been used by anyone else so far. We used uber-operon structures as the evolutionary footprints of cross-species, functionally related genes (**Fig. 1c**). We believe that genes are more likely to be functionally related if they have similar expression patterns and share the same uber-operon during evolution.

**Fig. 1.**
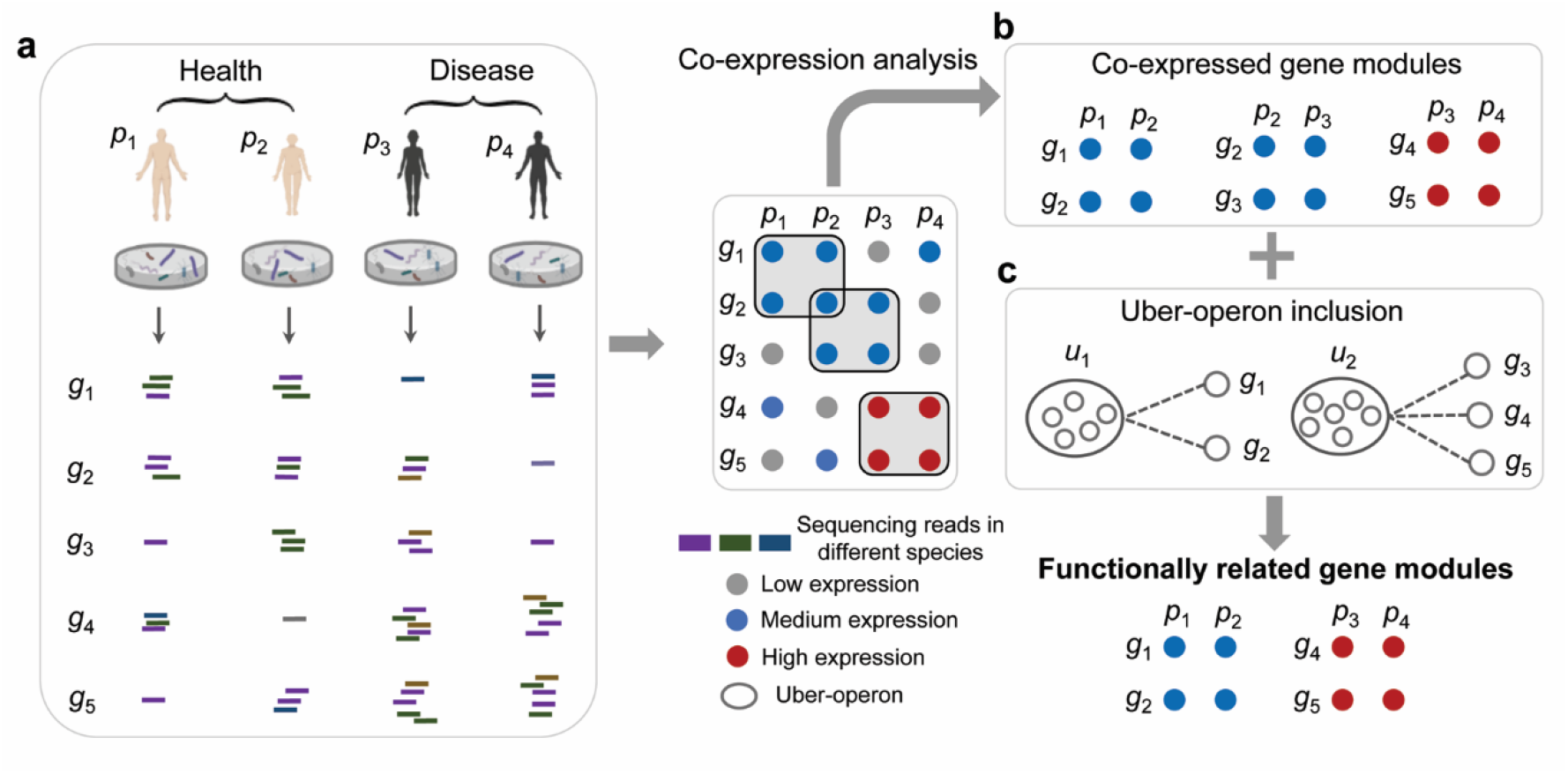
Key ideas of IDAM. **a**. Sequencing data from four samples (*p*_1_ and *p*_2_ from healthy cohorts, and *p*_3_ and *p*_4_ from disease cohorts). Since MG shotgun sequencing does not isolate microorganisms, the reads from different species (represented by purple, green, and blue) were jumbled together. We therefore could not distinguish them easily at the species level without alignment to reference genomes, but we could observe the community-level expression of each gene based on the merged reads from all species. The microbial community transcript number (MT data) can be corrected using the underlying genomic copy number (MG data) in practice. Here, we simply show the number of transcriptomic reads as expressions. The colors of the circles (grey, blue, and red) indicate the expression level of the genes within each sample. It is assumed that *g*_1_ and *g*_2_ are functionally related in healthy samples, while *g*_4_ and *g*_5_ are functionally related in disease samples. **b**. Co-expression modules. There were three co-expression modules, which are defined as genes with coordinated expression patterns under sample subsets. They were not equivalent to the known functional relationships. **c**. Gene modules by integrating uber-operon structures and expression similarity. Each ellipse, including multiple genes, represents an uber-operon. We searched for modules consisting of genes with both co-expressed patterns and shared uber-operons, which resulted in two identified biclusters. Therefore, we inferred there were functional-related relationships between *g*_1_ and *g*_2_, as well as *g*_4_ and *g*_5_, which were consistent with the ground truth.

We applied IDAM to published matched MG and MT datasets of inflammatory bowel diseases (IBD) (813 samples; including ulcerative colitis (UC) and Crohn’s disease (CD))^22^. The results suggested that IDAM performed better in key disease-associated characteristic inference than popular tools, including both supervised and unsupervised approaches. We further demonstrated the generalizability, reproducibility, and good performance of IDAM through independent cohorts with IBD and other publicly available datasets associated with melanoma^23^, type 1 diabetes mellitus (T1DM)^24^, and irritable bowel syndrome (IBS)^25^. Additionally, we found the obtained gene modules are likely to correspond to accessory genomic elements such as operons^15^. Hence, we believe that IDAM can be a highly advantageous method for disease-associated microbial characteristic inference based on MG and MT data. This approach will be valuable for investigating the relationships between microbiome and human diseases.

## Results

### IDAM: a metagenomic and metatranscriptomic analysis framework for disease-associated microbial gene module inference

We first present a formulation of the problem addressed by this article, and then provide an algorithmic overview of IDAM.

#### Computational formulation for gene module inference

The inference of gene modules can be mathematically formulated as an optimization problem. For matched MG and MT data, a gene-sample expression matrix *A*=(*a*_*ij*_)_*m*×*n*_ can be constructed, which includes *m* genes (*g*_1_,*g*_2_, ⃛,*g*_*m*_) and *n* samples (*p*_1_,*p*_2_, ⃛,*p*_*n*_). *a*_*ij*_ (1 ≤ *i* ≤*m*, 1 ≤ *j* ≤ *n*) represents the expression of gene within *g*_*i*_ sample *p*_*j*_, which is calculated by normalizing the RNA abundance (MT data) against the DNA abundance (MG data) of *g*_*i*_ within sample *p*_*j*_^26^. Based on the matrix, the co-expression patterns present in gene and sample subsets can be detected by identifying local low-rank submatrices^27^. Given the constraints of gene context conservation, we integrated the submatrices with a graph model to maximize the number of genes connecting to a single uber-operon. Specifically, we define a bipartite graph *G* = (*V*_1_,*V*_2_,*E*), where each node *u* ∈ *V*_1_ represents an uber-operon. Each node *v* ∈ *V*_2_ represents *a* gene in *A*, and the edge *e* ∈ *E* connecting a node *u* ∈ *V*_1_ to a node *v* ∈ *V*_2_ exists if the gene *v* belongs to uber-operon *u* (**Methods**). Using this graph and matrix *A*, the gene module inference problem can be formulated as the identification of a set of local low-rank submatrices within *A* that maximizes the number of connected components between the corresponding gene subsets and *V*_1_ on *G*. We maximize the number of connected components between gene subsets and *V*_1_, to minimize the heterogeneity of the gene subsets within the uber-operons, thereby ensuring that the genes within each identified submatrix are as functionally related as possible. Based on the identified submatrices within *A* (each contains a subset of genes and samples and we referred them to as biclusters hereinafter), the gene modules, i.e., the gene subsets, can be regarded as the characteristics of the shared phenotypes of the sample sets. Such an idea greatly alleviates the dependency on metadata in disease-associated characteristic inference. However, this problem is theoretically intractable (NP-hard, **Supplementary Information**). Hence, instead of solving this problem in a globally optimized way, we developed a heuristic algorithm, IDAM, as an approximate optimal solution.

#### Overview of IDAM

The main goal of IDAM is to identify disease-associated microbial gene modules (**Fig. 2a**). First, the community-level expression within each sample is assessed using HUMAnN2 based on matched MG and MT data^28^. This step provides a gene-sample expression matrix, each entry of which represents the expression of a gene within a sample. Second, IDAM combines the expression similarity and gene distribution within uber-operons to measure the likelihood that each pair of genes has the same function. This step provides a list based on the combined assessment for gene module initialization (**Fig. 2b**). Third, the gene pairs in the list are used as seeds, and each seed is expanded by iteratively adding other functionally related genes, to generate a bicluster until the new bicluster starts to become smaller than previously identified biclusters (**Fig. 2c**). Finally, the biclusters consisting of gene modules and corresponding sample sets are considered as the final output of the algorithm. We regard the gene modules as functional characteristics of the most common phenotype or state of the corresponding sample sets. Details are available in the Methods section.

**Fig. 2.**
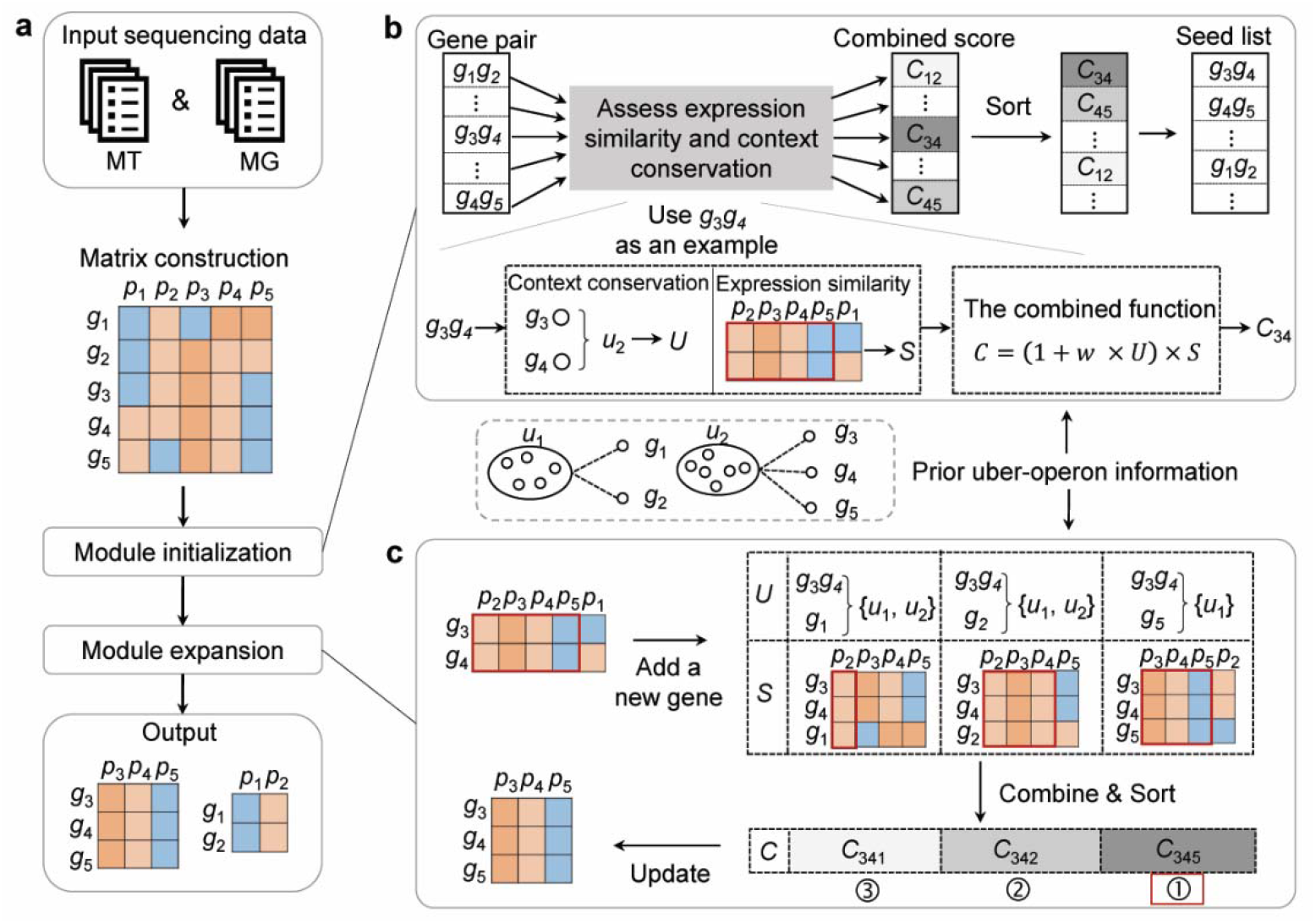
Workflow of IDAM. **a**. The flow chart. We consider the matched MG and MT data of multiple samples as the input, from which we obtain an expression matrix. By module initialization and expansion, the biclusters will be generated as the output. **b**. Module initialization. For each gene pair, we assess both expression similarity and context conservation to produce a combined score. Let us use *g*_3_*g*_4_ as an example. Based on known uber-operon structures, the two genes belong to *u*_2_. We assign a reward *U* for *g*_3_*g*_4_ as a context conservation measurement. The expression similarity is assessed according to the number of samples in which the rows are identical, and we obtain a score, *S* (Methods). Finally, *U* and *S* are combined via a function *C* (*w* is a tuning parameter), generating a score, *C*_34_. All gene pairs are sorted in a decreasing order based on the combined scores. **c**. Module expansion. We use gene pairs in the list and the samples under which the two genes are co-expression to obtain the initial biclusters and expand them gradually. We iteratively add a new gene with the highest combined score into a bicluster until the updated bicluster does not get bigger. Let us use *g*_3_*g*_4_ as an example. First, we add three genes (*g*_1_, *g*_2_, and *g*_5_) into *g*_3_*g*_4_, forming three new biclusters (denoted as *g*_3_*g*_4_*g*_1_, *g*_3_*g*_4_*g*_2_, and *g*_3_*g*_4_*g*_5_). Then, the combined score of each bicluster is assessed (**Methods**). Since bicluster *g*_3_*g*_4_*g*_5_ has the highest score, we replace the previous bicluster with this one. Next, we iteratively add a new gene (*g*_1_ and *g*_2_) to the current bicluster and implement the same processes. After comparing the combined scores, we find *g*_3_*g*_4_*g*_5_*g*_2_ to have the highest score, but the bicluster is of size 4 ×2, which is smaller than *g*_3_*g*_4_*g*_5_ (3 ×3). Therefore, we stop here and the bicluster *g*_3_*g*_4_*g*_5_ will be provided as the final output.

### The gene modules inferred by IDAM distinguished different phenotypes and enabled the discovery of disease subtypes

To assess the usefulness of IDAM, we assessed the phenotype classification performance of inferred gene modules. We applied IDAM to a publicly available dataset of patients with IBD and controls (referred to as non-IBD, **Methods**). The following analyses are based on the IBD-associated dataset unless otherwise specified. By assessing the community-level expression of each sample in the IBD-associated dataset, we obtained a matrix consisting of the expression of 917,127 genes across 734 samples. Based on this matrix, IDAM found 39 biclusters containing 47 to 5,452 genes (**Supplementary Table 1**), totally covering 29,127 genes. Ideally, samples with the same gene module should have highly consistent phenotypes. We used Z-tests to evaluate the phenotypic consistency of samples in each bicluster (**Methods**). Results suggested that 33 gene module associated sample sets were consistently enriched for a particular phenotype, with 20 enriched for IBD and 13 for non-IBD (**Fig. 3a and Supplementary Table 2**).

**Fig. 3.**
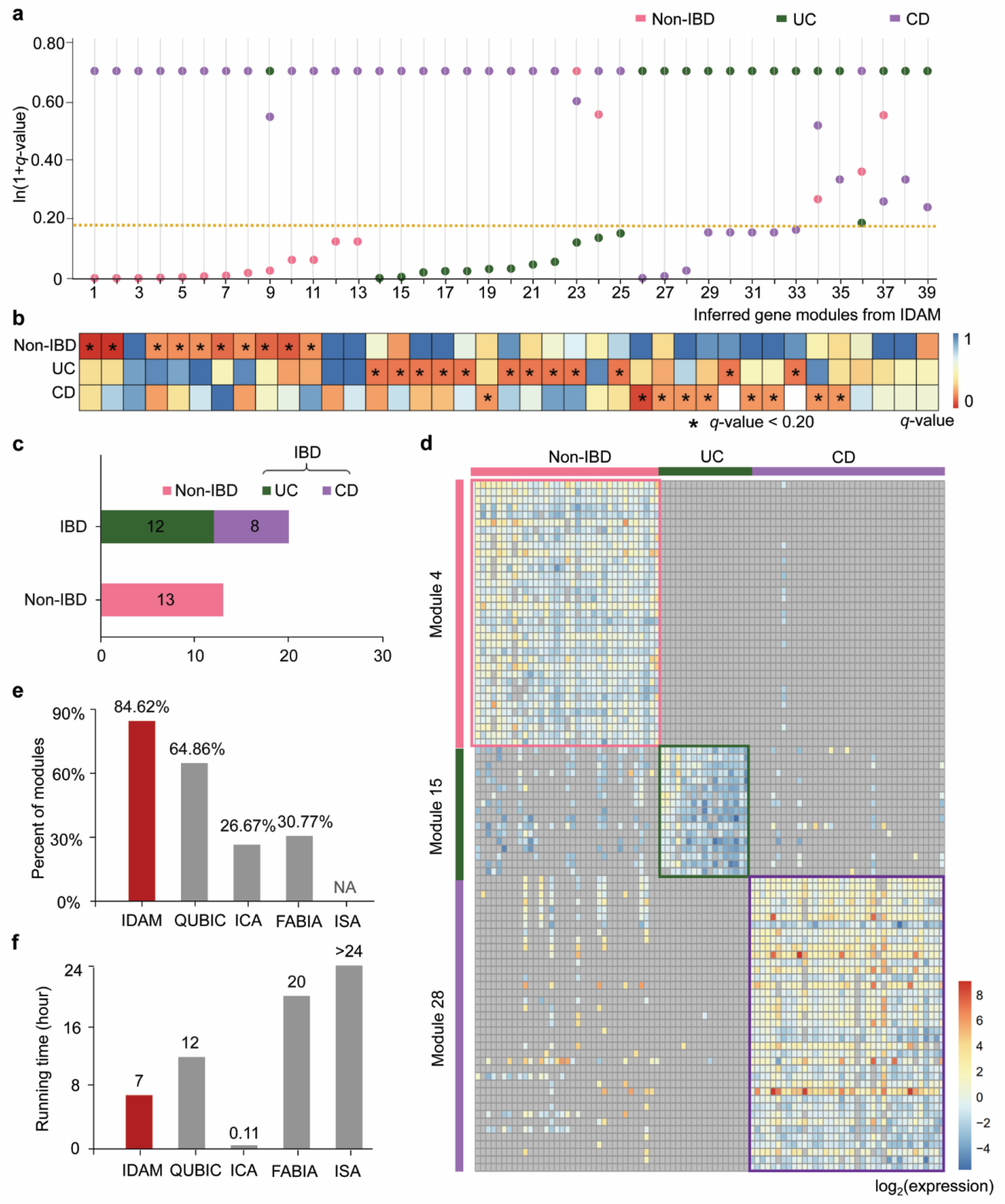
Performance of gene modules inferred by IDAM in distinguishing phenotypes. **a**. The values of ln(1+*q*-value) of non-IBD, UC, and CD, shown as pink, green, and purple, respectively. Each column represents a gene module. For modules 1-13 (except module 9), the ln(1+q-value) of UC and CD overlapped. For modules 14-22, the ln(1+q-value) of CD and non-IBD overlapped. We only showed the values corresponding to non-IBD in these cases. The yellow horizontal line indicates the critical value of *q*-value 0.20. The color of the circles below this line means that the corresponding phenotype exhibited strong enrichment within the gene module-associated sample sets. **b**. *q*-values of each gene module with respect to different phenotypes. Each column represents a gene module, and the color indicates the *q*-value. * indicates a *q*-value of < 0.20. **c**. The number of gene modules identified for each phenotype. **d**. An example of the gene modules associated with phenotypes. The color of each cell in the heatmap indicates the log2 value of the expression. Grey represents that the gene is not expressed within the corresponding sample. **e**. Percent of gene modules that were associated with samples enriched in certain phenotypes, identified by IDAM and competing algorithms. NA means no results from ISA. **f**. Running time of IDAM and competing algorithms.

We recognized that microbial communities can be influenced by multiple factors, such as age, race, and diet. These confounding factors can lead to high inter-individual heterogeneity, make disease-associated microbial characteristics less clear, and increase the risk of false positives or negatives in gene module inference. Hence, we further assessed whether the gene modules identified using IDAM had reliable associations with phenotypes, by applying a propensity score-based matching for each gene module to confirm its association with phenotypes. We constructed paired samples by matching the propensity scores estimated from other covariates, such as age, gender, and comorbidities. The Wilcoxon signed-rank test based on the paired samples indicated that 79.49% (31 of 39) gene modules were statistically significantly related to a specific phenotype, with *q*-values < 0.20 (**Fig. 3b and Supplementary Table 3**), demonstrating the high reliability of the gene modules identified using IDAM.

We specifically observed the enriched phenotypes of samples within each bicluster, and found 12 sample sets were with consistent phenotype UC and eight with CD, indicating the gene modules identified could recognize the subtypes of IBD (**Fig. 3c**). We here gave an example of three biclusters to show the phenotype specificity, namely Bicluster 4, Bicluster 15, and Bicluster 28 (**Fig. 3d**). Because of space limitations, we show the expression of the top 3% of genes in each bicluster: 35/1,162, 17/552, and 29/1264 for Biclusters 4, 15, and 28, respectively. IDAM detected the checker-board substructures from the original expression data. Gene module 4, corresponding to Bicluster 4, is specific to non-IBD, while Gene modules 15 and 28, corresponding to Biclusters 15 and 28, are specific to UC and CD, respectively, suggesting the capability of IDAM to uncover gene modules of disease subtypes. These gene modules were associated with 117 pathways (with *q*-values < 0.05 based on the hypergeometric test), of which 23 were also identified as differentially abundant across samples with different phenotypes based on pathway abundance from HUMAnN2 (**Methods and Supplementary Table 4**). For example, there was a significant enrichment in short-chain fatty acid metabolism (e.g., butyrate metabolism) in non-IBD-associated gene modules. The L-methionine biosynthesis, superpathway of L-citrulline metabolism, and superpathway of L-lysine, L-threonine, and L-methionine biosynthesis were enriched in non-IBD-associated gene modules, while preQ_0_ biosynthesis was enriched in IBD. The superpathway of lipopolysaccharide biosynthesis was enriched in CD-associated gene modules, and methylphosphonate degradation was enriched in UC-associated gene modules. These findings are in keeping with previous observations^29-32^, indicating that the gene modules from IDAM can provide functional insights into different phenotypes.

We also compared IDAM with several decomposition-based and biclustering-based approaches, including ICA^33^, QUBIC^34^, ISA^35^, and FABIA^36^. These methods have not been applied to microbial data for gene module inference, but they may reveal the functional characteristics of different phenotypes without the need for prior metadata. The results suggested that there were 37, 30, and 13 gene modules identified by QUBIC, ICA, and FABIA, respectively, while no gene module was detected by ISA. We further evaluated the phenotype consistency of gene module-associated sample sets using Z-tests. While 84.62% (33/39) sample sets from IDAM had a consistent phenotype, this number was only 64.86% (24/37), 26.67% (8/30), and 30.77% (4/13) for QUBIC, ICA, and FABIA, respectively (**Fig. 3e**). This finding indicated that the gene modules inferred by IDAM can cluster samples with the same phenotypes more effectively than other tools. This finding was not surprising, since IDAM took both gene expression and gene context conservation into consideration, thereby identifying more biologically meaningful gene modules, while others have used only gene expression. We also compared the running time of these methods (**Fig. 3f**). IDAM took seven hours, QUBIC and FABIA took 12 and 20 hours, respectively, while ISA was terminated after 55 hours without results. ICA took seven minutes; however, the application power of ICA was limited by the relatively low accuracy; only less than a third of the gene modules could distinguish between phenotypes. Collectively, the gene modules identified by IDAM performed best in distinguishing the phenotypes and can provide leads for new disease subtypes or states for future clinical testing.

### Species-level analysis suggested that the gene modules inferred by IDAM were biologically associated with phenotypes

Since differences in community-level functions ultimately imply species-level functional differences, we traced genes within each gene module back to individual species. These genes were from 133 species, which were dominated by Bacteroidetes and Firmicutes, followed by Proteobacteria at the phylum level (**Fig. 4a**). These were consistent with previous research into IBD-associated taxa^32^. To provide the genomic and functional context of the identified gene modules, we aligned the genes comprising gene modules to the reference genomes of these species in RefSeq^37^ and observed the distribution of genes on the reference genomes. We found the genes located on a genome tend to form several operons, instead of being spread out over the whole genome randomly. For example, the genes aligned to the representative genome of *Escherichia coli* make up several operons from RegulonDB^38^, such as operon *glnALG, ptsA*-*fsaB*-*gldA*, and *dcuB*-*fumB* (**Supplementary Fig. S2**). The operon *glnALG* encodes propanediol dehydratase, which has been demonstrated of higher levels in individuals with CD compared to that in healthy individuals^39^. This suggested the gene modules from IDAM were able to provide possible mechanisms for diseases that are suitable for further investigation.

**Fig. 4.**
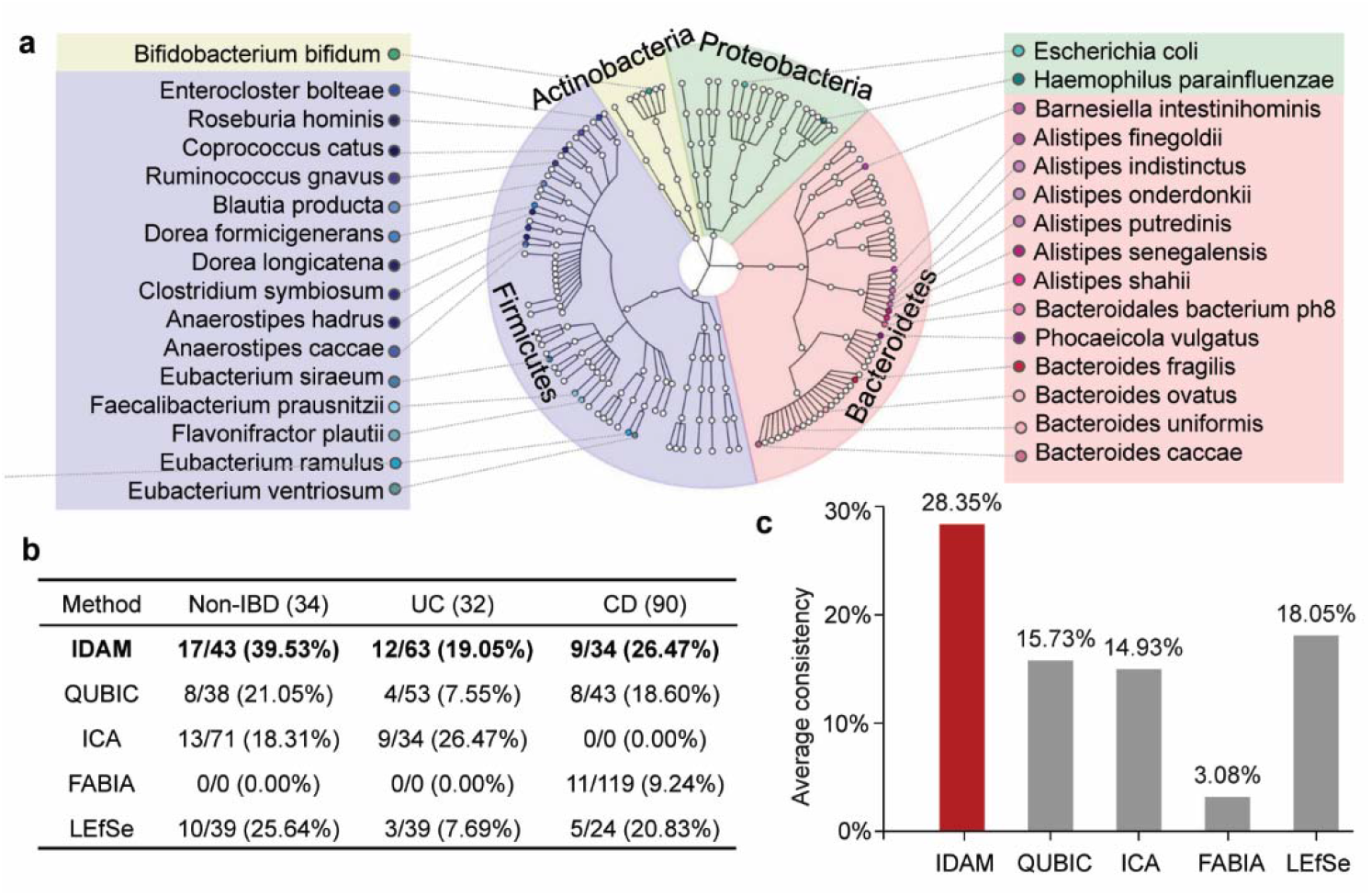
Species-level analysis of identified gene modules. **a**. Phylogenetic tree of the species associated with gene modules identified by IDAM. Colored dots indicate species that have been validated by previous studies. **b**. Results of species-level analysis by different methods. The numbers in the header (34, 32, and 90) represent the number of collected species associated with the three phenotypes, respectively. Each cell in the table is represented in the form of X/Y (Z). 1) The denominator Y means the number of species associated with a particular phenotype identified from the different methods; 2) X means the number of identified species that are consistent with the collected species; and 3) Z indicates the consistency index of each method associated with the phenotypes, which reflects the reliability of identified species of different methods. **c**. Average consistency for each method. A higher value indicates that inferred species are more reliable.

We further classified the species according to gene module-associated sample phenotypes from Z-tests and obtained 43, 63, and 34 species associated with non-IBD, UC, and CD, respectively (**Methods and Supplementary Table 5**). Seven of these species overlapped between UC and CD. We collected IBD-related species (34 for non-IBD, 32 for UC, and 90 for CD) from Peryton, a database of microbe-disease associations, and previous taxonomic studies into IBD^40-43^ (**Supplementary Table 6**). Among them, a total of 31 species were captured by IDAM (**Fig. 4a**), including 17 for non-IBD, 12 for UC, and 9 for CD (shown in red in Supplementary Table 6). The seven overlapped species from IDAM have been reported to be associated with both UC and CD^40-42^, suggesting that the IDAM results are reliable. We compared IDAM with QUBIC^34^, ICA^33^, FABIA^36^, and LEfSe^11^. ICA, QUBIC, and FABIA are not dependent on metadata, while LEfSe identifies differentially abundant species based on prior phenotype information. Detailed results are shown in **Fig. 4b**. The average consistency with collected species across the three phenotypes was higher for IDAM (28.35%), than for QUBIC (15.73%), LEfSe (18.05%), ICA (14.93%), or FABIA (3.08%) (**Fig. 4c**). The species-level analysis showed that the gene modules inferred by IDAM were biologically associated with phenotypes.

### Integration of gene context conservation substantially improved the performance of gene module inference

Our hypothesis is that genes with similar expression patterns and high context conservation during evolution are likely to be functionally related. To evaluate the validity of this hypothesis, we sought to identify gene modules without using gene context conservation in IDAM (referred to as IDAM without gene context conservation). As a result, we found the gene modules identified by this version were less capable of distinguishing different phenotypes than those from IDAM. IDAM without gene context conservation detected 37 biclusters from the expression matrix, of which only 64.86% (24/37) sample subsets were of consistent phenotypes (the percentage of IDAM is 84.62%). Pathway analysis suggested that 78.38% (29/37) gene modules from IDAM without gene context conservation were significantly associated with MetaCyc functional categories (with *q*-values < 0.05 using the hypergeometric test), which is lower than the 94.87% (37/39) of IDAM (**Fig. 5a**). These findings indicated that the integration of uber-operon structures contributed to the identification of informative gene modules.

**Fig. 5.**
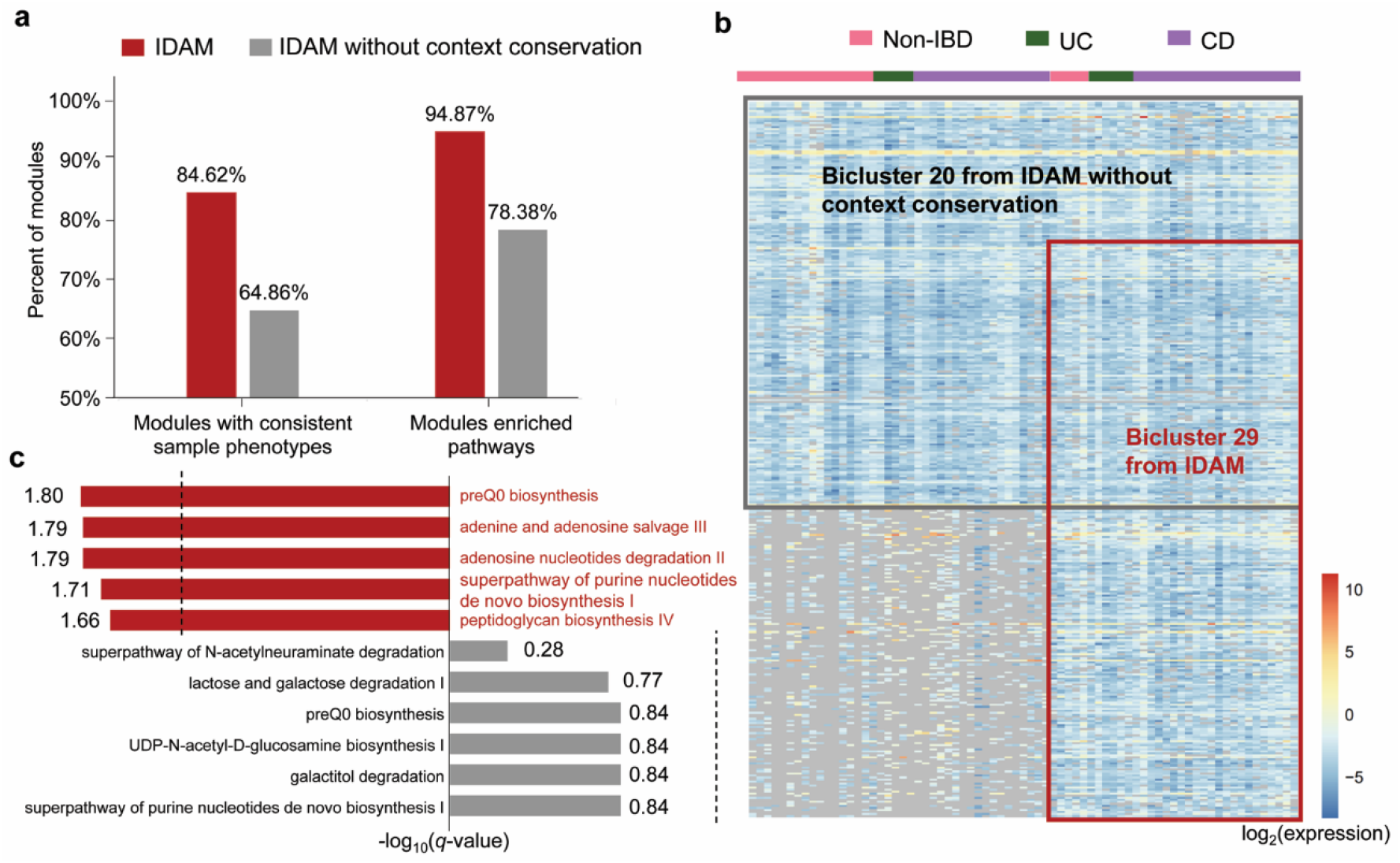
Analysis of the contribution of gene context conservation. **a**. Comparison between IDAM and IDAM without gene context conservation. **b**. A specific case of identified biclusters. Each row represents a gene, and each column represents a sample (a total of 474 genes and 73 samples). The colored bar at the top represents the phenotypes of the corresponding samples. **c**. Pathways enriched in Bicluster 20 and Bicluster 29. The pathways in grey indicate the pathways enriched in Bicluster 20, while red indicates those enriched in Bicluster 29. The dotted vertical lines indicate the critical value (a *q*-value of 0.05). The bars located outside the dotted lines correspond to significantly enriched pathways.

We explored the underlying reasons for the superior performance of integrating gene context conservation. We observed the distribution of the genes in the uber-operons for each gene module from both IDAM and IDAM without gene context conservation. Overall, the genes within gene modules that are not significantly associated with any pathways tended to be spread out among more uber-operons. This finding supports our hypothesis that functionally related genes are of high context conservation. As an example, we investigated Bicluster 20 from IDAM without gene context conservation (abbreviated as ‘Bicluster 20’ hereinafter, **Fig. 5b**). Since the 200 genes within the bicluster had coordinated expression across the 73 samples, they would be clustered together if we merely considered co-expression. However, these genes were not significantly functionally related, and were from eight uber-operons. They cannot reflect the unique functional characteristics of a particular phenotype. Indeed, the 73 samples within Bicluster 20 were evenly scattered over different phenotypes: 23 in non-IBD, 11 in UC, and 39 in CD. When we took gene context conservation into consideration, a new bicluster occurred (Bicluster 29 from IDAM, abbreviated as ‘Bicluster 29’ hereinafter).

This bicluster consists of 328 genes and 33 samples, and there was considerable overlap with Bicluster 20 (**Fig. 5b**). The bicluster was dominated by an uber-operon that is closely associated with purine metabolism^21^, and the enriched pathways are shown in **Fig. 5c**. Previous studies have shown that the imbalance between the biosynthesis and degradation of purines can produce excessive uric acid in the gut, leading to the occurrence of IBD^44^. The facts showed the 33 samples indeed were significantly enriched in phenotype CD, with a *q*-value < 0.20 using a Z-test. This finding demonstrated the value of gene context conservation in the clustering of genes with biological relevance, which improves the ability of IDAM for gene module inference.

### IDAM showed high reproducibility and good generalizability in gene module inference and provided insights into melanoma-related progression-free survival after immunotherapy

We used an independent dataset associated with IBD to validate the reproducibility of inferred gene modules (**Methods**). For each gene module associated with IBD inferred from IDAM, we estimated the mean difference in centered log-ratio expression between samples with and without certain phenotypes using the independent dataset. The random effects, such as age and diet, were balanced by matching the propensity score of the samples. Results suggested the expression of 79.49% (31/39) gene modules was significantly different across phenotypes, with a *q*-value < 0.20 (**Supplementary Fig. S3a**). That means these gene modules were also significantly associated with sample phenotypes on the independent dataset. When performing the same analysis on single gene expression (a single gene randomly selected from each gene module), we observed a much lower rate (33.33%) with significant differences across phenotypes (with *q*-values < 0.20, **Supplementary Fig. S3b**). This supported the utility of gene modules for exploring reproducible associations between microbiome and diseases.

To demonstrate the generalizability of IDAM, we applied it to an integrated dataset containing 60 samples from four different cohorts: cohorts with melanoma, T1DM, without T1DM (non-T1DM), and IBS (**Supplementary Information and Supplementary Table 7**)^23-25^. The data processing included quality control and the measurement of the community-level expression of each sample (**Supplementary Information**). A matrix with the expression of 470,305 genes across 60 samples was produced. Based on this matrix, IDAM inferred 55 gene modules covering 13,145 genes. Analogous to the analyses of the IBD-associated dataset, we first evaluated the phenotype consistency of each gene module-associated sample set using Z-tests and found that 32 of 55 (58.15%) were consistently enriched for a particular phenotype: seven enriched for T1DM, 18 for non-T1DM, and seven for melanoma. Comparison with other methods, QUBIC, ICA, FABIA, and ISA, showed that the gene modules inferred by IDAM could distinguish between the phenotypes most effectively (**Fig. 6a and Supplementary Table 8**). We then found that 20.43% of the species from IDAM were consistent with the collected disease-specific species, and the average consistency across the four phenotypes was the highest of all the methods (**Fig. 6b and Supplementary Table 9**). Pathway enrichment for the genes within each gene module showed that IDAM could detect eight differentially abundant pathways identified by HUMAnN2 (**Supplementary Table 10**) and many pathways consistent with previous findings^45-47^. For example, the biosynthesis of low-carbon-number saturated lipid classes, including myristic, stearic, and palmitic acid, was likely to be higher in those with T1DM than in those without T1DM^46^. Aromatic amino acids, phenylalanine, and tyrosine were higher in T1DM patients^45^, which tends to increase the risk of T1DM. Collectively, these results demonstrated the high reliability of the gene modules identified by IDAM.

**Fig. 6.**
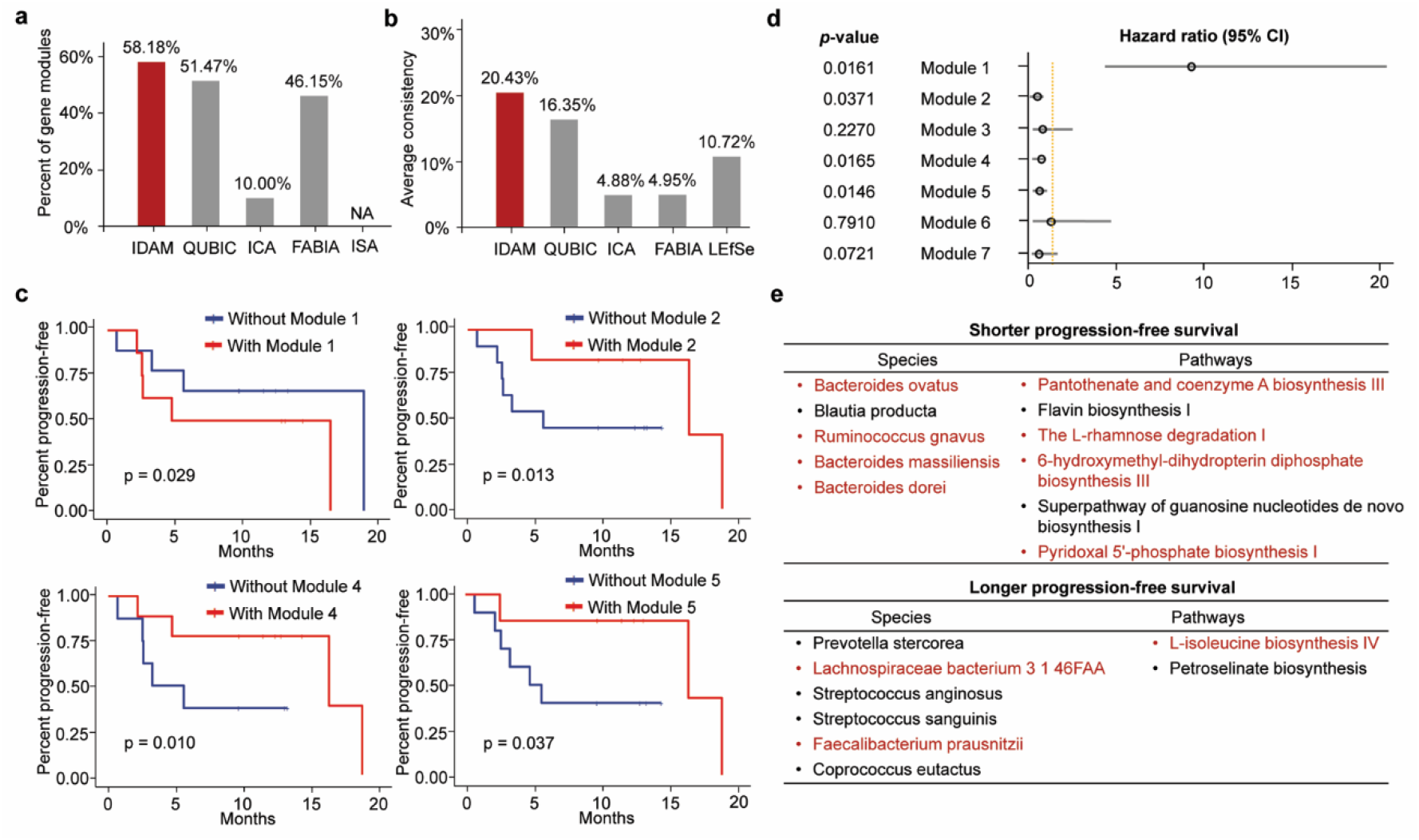
Results of IDAM on the integrated dataset. **a**. Comparison of phenotype consistency among IDAM, QUBIC, ICA, FABIA, and ISA. The red bars indicate the percent of gene module-associated sample sets with consistent phenotypes from IDAM, while the grey bars indicate those from other methods. **b**. Comparison of the average consistency of collected species across the four phenotypes. **c**. Kaplan-Meier curves of gene module-associated groups along with corresponding log-rank test *p*-values. **d**. Results from Cox proportional hazard models. The yellow line indicates a hazard ratio equal to 1. CI: confidence interval. the *p*-value is from the likelihood-ratio test. **e**. Species and pathways related to PFS. The red highlights represent those that can be found by gene modules from IDAM.

We also noted that progression-free survival (PFS) after immunotherapy of melanoma individuals was publicly available. We explored the association between inferred microbial gene modules of melanoma and PFS (**Methods**). Four out of seven gene module-related sample groups, with or without the gene module, differed significantly in their PFS, with a log-rank *p*-value < 0.05 (**Fig. 6c and Supplementary Fig. S3c**). We assessed the progression risk of patients using Cox proportional hazards models, and found that the patients with Gene modules 2, 4, and 5 had a lower risk of progression than those without these gene modules (**Fig. 6d and Supplementary Table 11**), as assessed during follow-up. Thus, we inferred that the three gene modules were associated with longer PFS. In contrast, gene module 1 was associated with shorter PFS, and individuals with this gene module had a high risk of melanoma progression (Hazard ratio [95% CI] = 9.1195 [3.987,20.615]; **Fig. 6d**). We collected all species and pathways that have been identified to be associated with shorter or longer PFS from the paper by Peters *et al*.^23^ (**Fig. 6e**). Most of these (6/11 species and 5/8 pathways) can be found in the gene modules identified by IDAM (shown as red in Fig. 6e). These findings support the contention that the gut microbiome may have an impact on the efficiency of immunotherapy on melanoma^23^, and provide possible mechanistic insights into means for improving the efficiency of immunotherapy by manipulating the microbiome.

## Discussion

Although many studies have focused on the microbial characteristic inference of diseases^9-13^, it is still a fundamental challenge to explore critical characteristics in large-scale, highly heterogeneous microbial data. We developed a new methodology, IDAM, for matched MG and MT data to identify disease-associated gene modules. The above results indicate that IDAM is a highly effective approach to the identification of disease-associated microbial gene modules based on both gene co-expression and context conservation during evolution. The gene modules of diseases identified can be useful in understanding disease pathogenesis and providing guidance for the future development of disease treatment.

IDAM is innovative in that, unlike previous disease-associated microbial characteristics analysis methods^9-13^, it integrates gene context conservation and gene expression information by leveraging a multi-omic view of microbial communities for the inference of diseases using functional characteristics. IDAM does not rely on prior knowledge about the samples. Given that existing metadata is often misannotated and misleading, even publicly unavailable^18^, creating an overly complex environment for sample reanalysis. This will reduce the application power of previous metadata-based methods. IDAM can be a good complement to those methods in this case and eliminates bias from misleading data noise. Additionally, IDAM, as an unsupervised method, is effective to develop new biological hypotheses, providing novel insights into new disease subtypes or stages. Previous supervised learning methods, by contrast, cannot provide such information because they identify key features based on known labels and annotations of samples.

We illustrated and evaluated IDAM using IBD, melanoma, T1DM, and IBS associated datasets, and found that it outperformed existing methods. We discovered microbial gene modules significantly associated with specific diseases, some of whose functions are consistent with what has been reported previously. We also evaluated the gain of using gene context conservation information in IDAM and found that ignoring this information can lead to significantly decreased biological relevance between genes within a bicluster. This research demonstrated that uber-operon structures play a crucial role in functional indication. Furthermore, IDAM is suitable for the inference of microbial characteristics on large-scale datasets. One can integrate microbial datasets spanning multiple diseases and include multiple independent studies for each disease to explore disease-specific microbial characteristics^48^, providing possible insights into disease diagnosis.

IDAM is still not free of limitations. It identifies gene modules at the community level, which need further taxonomic analysis to improve their clinical utility. This limitation can be somewhat addressed by tracing genes back to taxa by mapping them to existing reference genomes. However, there are some genes that cannot be attributed to known species, which would lead to the biological information in the gene module being inadequately interpreted. We believe that this issue will be alleviated as more microbial species and reliable metagenome-assembled genomes are discovered and explored. We plan to develop a three-dimensional analysis approach in the future^49^, which takes both microbial species and genes and pathways of multiple samples into consideration for disease-associated microbial characteristic inference. In this way, modules involved in diseases consisting of genes and pathways with specific functions of specific species will be detected, providing both functional and taxonomic insights into diseases. Additionally, IDAM requires relatively large amounts of data. Since IDAM uses expression similarity to assess the relevance of gene regulation, a large sample size might be needed to guarantee reliable measurement from data. In practice, new paired MG and MT data from users can be integrated with the datasets used in this article to increase the sample pool, thereby improving the practicability of IDAM. Finally, the gene modules identified by IDAM only provided associations between microbiomes and diseases. Rigorous biological experiments are required to establish the causality between the gene modules identified and the disease of interest, and to establish their clinical prevalence and utility.

## Methods

### Datasets

We downloaded all matched quality-controlled MG and MT data associated with IBD, as well as the corresponding HUMAnN2 results and metadata of samples at the IBDMDB website in March 2020 (https://ibdmdb.org)^22^. There were 198 samples from non-IBD subjects and 615 samples from patients with IBD, consisting of 381 UC samples and 234 CD samples. These samples were collected in two phases: an initial round as a pilot obtaining 79 samples with matched MG and MT data and the second phase obtaining 734 samples with matched MG and MT^22^. We split the IBD-associated dataset into two parts based on sample collection phases: one consisting of 79 samples used for validation and the other consisting of 734 samples used for gene module inference. We also downloaded matched MG and MT data in .fasta format from the NCBI Sequence Read Archive with the accession number of SRP197281 for melanoma-associated data^23^, SRP072561 for T1DM- and non-T1DM-associated data^24^, and ERP001739 for IBS-associated data^25^. Uber-operon structures were obtained from the paper by Che *et al*.^21^. The authors predicted the uber-operons through identifying groups of functionally or transcriptionally related operons whose gene sets are conserved across multiple reference genomes^21^.

### IDAM: a framework for identifying disease-associated microbial gene modules based on metagenomic and metatranscriptomic data

IDAM is a heuristic algorithm for disease-associated microbial gene module inference. It is based on a greedy search to gather genes with similar expression patterns and high context conservation as functional gene modules. IDAM consists of three steps.

### Step 1: Extracting community-level gene expression of samples

The input data is quality-controlled (quality- and length-filtered and screened for residual host DNA) matched MG and MT sequencing datasets in fasta, fasta.gz, fastq, or fastq.gz format. The datasets of each sample are first processed using HUMAnN2 for gene relative expression within communities^28^. Here, we performed both nucleotide-level search and accelerated translated search using HUMAnN2 for a more comprehensive community-level profiling than only reference genomes based mapping^28^. Given that random sampling may lead to the detection of transcripts for undetected genes for low-abundance species, we used Laplace smooth for RNA and DNA abundance and use smoothing DNA-level features to avoid divide-by-zero errors during normalization. In this study, we directly used the HUMAnN2 results of the datasets from IBDMDB for the IBD-associated datasets. Then, we extract the community-level gene expression of each sample to construct a gene-sample expression matrix *A*_*m* ×*n*_, in which each row represents a gene (total *m* genes indicating by UniRef90 identifiers^50^ and forming a gene set *X*), and each column represents a sample (total *n* samples).

### Step 2: Measuring gene context conservation and expression similarity for each gene pair

We assess context conservation and expression similarity for each gene pair and sort them as a list for module initialization.

#### 2.1 Gene context conservation

Gene context conservation is assessed based on the gene distribution in uber-operons. Since the genes within *X* referred to here are indicated as gene families from UniRef90 (a non-redundant protein sequence database)^50^, we align the gene family sequences with genes in uber-operons to determine to which uber-operons these gene families belong. The sequences of genes in all uber-operons are collected into a custom database. For each gene family, *g*_*i*_, within *X* (1 ≤ *i* ≤ *m*), we extract the corresponding sequence from the UniProt database^50^ and align it with the custom database using BLAST^51^. We define a gene-to-uber-operon mapping function φ *D* → *H*, in which *D* consists of all subsets of *X, Y* is the set of all uber-operons, and *H* consists of all subsets of *Y*. We assume that a gene family *g*_*i*_ (1 ≤ *i* ≤ *m*) belongs to an uber-operon *u* if the *E-*value of its alignment with one of the uber-operon’s sequences is less than 0.001. This is denoted by φ(*G*_*i*_) = {*u*}, where *G*_*i*_ is the set consisting of a single gene, *g*_*i*_. If multiple sequences from different uber-operons have an *E-*value less than 0.001, we choose the uber-operon corresponding to the smallest *E*-value. In contrast, if no sequence has an *E-*value less than 0.001, we set φ(*G*_*i*_) = {ø}. In this way, we construct a map between an uber-operon and a gene.

We further define the relationship between uber-operons and gene modules with multiple genes. Suppose a gene set *Q* consists of *b* genes {*g*_1_,*g*_2_, ⃛,*g*_*b*_} (1 < *b* ≤ *m*). We assume φ(*Q*) = φ ({*g*_1_} ∪ {*g*_2_} ∪ ⃛ ∪{*g*_*b*_}) = φ(*G*_1_ ∪ *G*_2_ ∪ ⃛ ∪ *G*_*b*_) = φ(*G*_1_) ∪ φ (*G*_2_) ∪ ⃛ ∪ φ (*G*_*b*_), where *G*_*i*_ represents the set consisting of a single gene *g*_*i*_ (*i* = 1,2, ⃛, *b*). For two gene modules, *I* and *J*, we set a reward *U*(*I,J*) as a gene context conservation measurement, based on the relative positions of their genes in uber-operons:

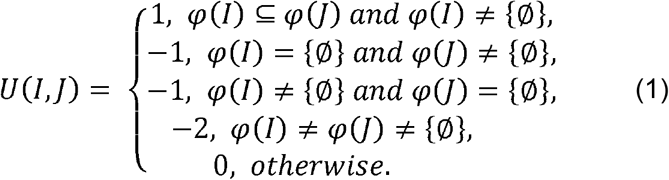

Based on this, we calculate *U* (*G*_*i*_, *G*_*j*_) as the gene context conservation of each pair of genes *g*_*i*_ and *g*_*j*_ (1 ≤ *i* ≤ *j* ≤ *m*).

#### 2.2 Expression similarity of genes

Expression similarity is assessed based on a vector composed of the expression values of each gene within all samples. We need to measure the expression similarity not only between two genes, but also across multiple genes, while expanding the modules in *Step 3*. This cannot be achieved using existing similarity measurements, such as Pearson correlation coefficients, since they are generally used for paired data points. To address this issue, the continuous expression value of each gene is first discretized to an integer representation using qualitative representation^34^ (**Supplementary Information**). Based on this representation, we measure the expression similarity *S* (*G*_*i*_, *G*_*j*_) of gene pair (*g*_*i*_, *g*_*j*_) as the number of samples under which the corresponding integers along the rows of the two genes are identical (or identical integers but with opposite signs).

#### 2.3 Integration of gene context conservation and expression similarity

A combined score, *C* (*G*_*i*_,*G*_*j*_), integrating gene context conservation and the expression similarity of *g*_*i*_ and *g*_*i*_ is obtained via the following function:

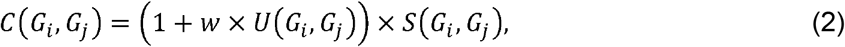

where *w* is a tuning parameter that can be set by users. *w* ranges between 0 and 1, and a larger value of *w* indicates that gene context conservation has a stronger impact on estimating functional relationships. Here we used 0.3 as the default value (**Supplementary Information and Supplementary Table 12**). Finally, the gene pairs are sorted in descending order for the combined scores, generating a seed list, *L*.

*Step 3*: *Generating biclusters consisting of gene modules and corresponding sample sets* Since the gene set *X* is a pan-genome of all microorganisms from multiple samples, it contains a large number of genes. To improve the computational efficiency, we partition the rows within the matrix equally into *t* subsets^52,53^, where *t* is determined using a stochastic model^52,53^. Then, we use gene pairs within each subset instead of all gene pairs in *X*, forming the seed list *L*, which can greatly decrease the number of seeds and reduce the computational burden.

A submatrix is regarded as biologically relevant if it does not occur randomly in a matrix *A*_*m*×*n*_. Since the output submatrices from our algorithm are biologically relevant and generated by a gene pair (see *Step* 3.1), we need to make sure that at least one gene pair within each meaningful submatrix can exist as a seed, to keep all biologically relevant submatrices detectable. This means we can at most divide the rows of matrix *A* into *k* - 1 subsets (*i*.*e*., *t= k* −1), where *k* represents the number of genes of the biologically relevant submatrix with the fewest genes (referred to with the smallest row size). We statistically assess the smallest row size of the submatrix that is unlikely to occur by chance. The submatrices we target are of rank one mathematically, since the corresponding genes have co-expressed patterns. Suppose a submatrix with rank one is of size *k*×*s*, and its probability of random occurrence in *A*_*m*×*n*_ is calculated as follows^52,53^:

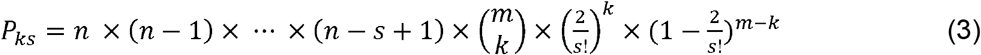

The submatrix is considered to be biologically relevant if *P*_*ks*_ < 0.05. When users have prior knowledge of the largest column size, *s*, we can calculate *k* based on the formula above. The smallest row size of the biologically relevant submatrices can be determined.

Users can then set the number of subsets (*t= k* −1). In this work, the default value of *t* was set to 10. That is, the matrix was divided into 10 subsets, which greatly reduced the computational burden and kept biologically relevant submatrices detectable to a large extent.

#### 3.1 Initialization

We start with the first gene pair in *L* that satisfies one of two conditions: i) at least one gene of these two has not been included in previous biclusters, or ii) these two genes are included within different previous biclusters, and there is no overlap between the gene components of the two biclusters. We aggregate samples in which the integers of the two genes are identical (represented as set *P*), forming a current bicluster *M=* {*Q, P*}, where *Q* represents the gene set consisting of the paired genes. The gene pair is then removed from *L*.

#### 3.2 Expansion

We expand *M* by adding a new gene (if any) that has the highest combined score with *M*, giving rise to the updated bicluster *M*′ *=* {*Q*′, *P*′}. The calculation of the combined score between a gene (represented as *g*_*z*_ (1 ≤ *z* ≤ *m*)) and *Q*, namely *c*(*G*_*z*_, *Q)*, is similar to that of two genes *g*_*i*_ and *g*_*j*_. We first assess the context conservation between *g*_*z*_ and the gene set *Q* of bicluster *M*, i.e., *U*(*G*_*z*_, *Q)*. Then, we measure the similarity of expression of *g*_*z*_ and *Q, S*(*G*_*z*_, *Q)*, as the number of samples within *P* under which the corresponding integers along the rows are identical (or identical integers but with opposite signs). Based on these, a combined score *c*(_*Gz*_, *Q)* can be obtained by replacing *U*(*G*_*i*_, *G*_*j*_) and *s* (*G*_*i*_, *G*_*j*_) in the calculation of *c* (*G*_*i*_, *G*_*j*_ with *U*(*G*_*z*_, *Q)* and *S*(*G*_*z*_, *Q)*, respectively. If *min* |*Q*′|, |*P*′|} *> min* |*Q*|, |*P*|}, we set *M= M*′ and repeat Step 3.2. Otherwise, go to *Step* 3.3.

#### 3.3. Filter and output

If the proportion of genes overlapping between *Q* and the component genes of previously identified biclusters is less than a threshold, *f*, bicluster *M* will be provided as the final output, and is ignored otherwise. In this study, the default *f* was set to 0.1 for more separate biclusters. Go to *Step* 3.1 until the list *L* is empty.

##### Evaluating phenotype consistency of samples within one bicluster

Z-tests were used to assess whether samples with the same gene module are of highly consistent phenotypes. Here, the samples in each bicluster were regarded as with the corresponding gene module. We extracted the phenotypes of all samples from the metadata as background, and the phenotypes of samples in each bicluster as test sets. For each phenotype and each test set, we counted the phenotype occurrence in the test set and background. We evaluated the significance of each phenotype using a right-tailed Z-test with the null hypothesis that the observed frequency in the test set is less than or equal to the occurrence probability within the background. The resulting Z-score is tranfered to *p*-value, and further adjusted using the Benjamini-Hochberg procedure for false discovery rate. If the *q*-value is less than 0.20, the null hypothesis is rejected, meaning the phenotype significantly occurred within the test set. In this case, we consider the samples within the bicluster to be of a highly consistent phenotype.

##### The propensity score matching method

Our aim was to assess whether the association between identified gene modules and disease phenotypes was reliable. For each gene module, we first identified the underlying confounding variables that affected its occurrence. We included diet, age, race, and health status, as well as the other gene modules, and calculated a propensity score for each sample using the R package MatchIt 4.2.0. We then matched each sample with the gene module to a sample without the gene module but has a similar propensity score with the former, obtaining paired samples consisting of samples with and without the gene module. This was achieved by setting the parameter method = “optimal”. Finally, for each phenotype, we used Wilcoxon signed-rank test to test the null hypothesis that there is no difference in the phenotype occurrence between sample groups with the gene module and those without. The *p*-value was adjusted using the R package qvalue 2.18.0. The *q*-value less than 0.20 means the gene module is significantly associated with the phenotype.

##### Functional analysis of gene modules

We performed an analysis of enriched functions for each gene module. All UniRef90 identifiers were firstly mapped to MetaCyc reactions^28,54^. The reactions were further associated with MetaCyc pathways^28,54^, which enabled a transitive association between UniRef90 gene families and MetaCyc pathways. We defined a pathway that was satisfied when the reactions within the pathway significantly occurred via the hypergeometric test,

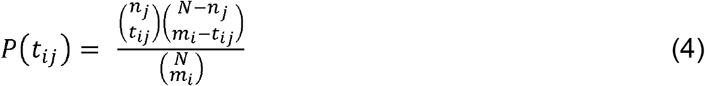

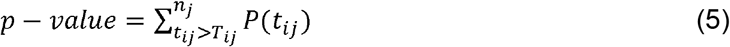

where *N* is the total number of reactions that were associated with the UniRef90 gene families and pathways in MetaCyc, *m*_*i*_ is the number of reactions within the *i*th gene modules, *n*_*j*_ is the total number of reactions within the th pathway, and *T*_*ij*_ is the number of observed reactions of pathway *j* occurring in gene module *i*. This yielded a *p*-value for each gene module and pathway. The resulting *p*-values were adjusted using the Benjamini-Hochberg procedure for false discovery rates. We selected the pathways with a *q*-value less than 0.05 as the significantly enriched pathways of each gene module.

We used a widely used tool, LEfSe^11^, to infer differentially abundant pathways among different phenotypes. The pathway abundance information for each sample is from HUMAnN2. Then, we compared the differentially abundant pathways and significantly enriched pathways of gene modules from IDAM.

##### Comparison of IDAM with other methods

The performance of gene module inference of IDAM was compared with the other four top-performing unsupervised tools for grouping genes into functional modules ^55^, including biclustering-based (QUBIC^34^, ISA^35^, and FABIA^36^) and decomposition-based (fastICA, an improvement of ICA^33^) methods. All tools were tested and compared based on the gene-sample matrix constructed. We ran these tools on the Pitzer cluster of the Ohio Supercomputer Center with memory usage set to 300GB^56^ with default parameters. QUBIC was implemented using the published source code, while the other three algorithms were implemented using the R packages, isa2 0.3.5, fabia 2.36.0, and fastICA 1.2.2, respectively.

##### Species-level analysis

For the gene modules from IDAM, ICA^33^, QUBIC^34^, and FABIA^36^, we extracted species that contribute to the abundance of genes within the identified gene modules, based on the HUMAnN2 output^28^. In this way, community-level gene expression (abundance) was decomposed into species-level data. A phylogenetic tree was generated using the software GraPhlAn^57^. Since the gene modules have been associated with phenotypes using Z-tests, we assigned the species of each gene module to the corresponding shared sample phenotype of the samples within the same bicluster. We collected disease-associated species from the databases Peryton and MicrophenoDB^58^, and previous taxonomic studies. For each phenotype, we counted the total number of species, as well as the number that matched with the collected species. Then, the consistency index was calculated as the total number of species in the phenotype dividing by the number of matched species. We used the average of the consistency indexes across the three phenotypes as the average consistency index.

LEfSe^11^, was used here for comparison. It uses species profiling from MetaPhlAn2 to identify statistically different species among phenotypes^59,60^. The significantly different species of each phenotype were aligned to the collected species, from which we determined the number of matched species and calculated the consistency of each phenotype.

##### Gene module validation

We used the 79 samples from an initial round associated with IBD as an independent dataset to validate the reproducibility of gene modules from IDAM. For each gene module inferred from IDAM (a total of 39 modules), we test the null hypothesis that the mean difference in centered log-ratio expression between patients with and without certain phenotypes was zero using Wilcoxon signed-rank test based on propensity score-based paired samples. We calculated the *q*-values using the qvalue R package 2.18.0. The *q*-value less than 0.20 means there was a significant difference in the mean expression of the gene module between patients with and without the phenotype. That was, the gene module was significantly associated with the phenotype of samples in the independent dataset.

##### Survival analysis

We assessed the associations of the seven identified gene modules of melanoma with PFS, respectively. The samples in each melanoma-associated bicluster were considered as with the corresponding gene module. In this way, we determined which gene modules a sample contains. Given that a sample may contain multiple gene modules, it may be influenced by other gene modules when measuring whether a gene module is associated with PFS. For each gene module, we first took all the other gene modules as underlying confounding variables to calculate the propensity score for each sample and constructed paired sample groups, corresponding to with or without the gene module groups. Then, the Kaplan-Meier curves of each group were plotted using the “ggsurvplot” function in the survival R package^61^. The Cox proportional hazards model was used to explore specific relationships between the gene module and PFS, adjusting for age, antibiotic use in the last 6 months, sex, BMI, and disease stage. This was achieved using the “coxph” function in the survival and survminer R packages^62^. Additionally, we identified species and pathways associated with the gene module with shorter (Gene module 1) or longer PFS (Gene modules 2, 4, and 5), which is the same as the description in subsections “Functional analysis of gene modules” and “Species-level analysis”.

## Data availability

The IBD-associated dataset supporting the conclusions of this article is available in the IBDMDB database (https://ibdmdb.org). The melanoma, T1DM, and IBS associated datasets supporting the conclusions of this article are available on NCBI Sequence Read Archive, with the accession number of SRP197281, SRP072561, and ERP001739, respectively.

## Acknowledgements

This work was supported by National Key R&D Program of China (2020YFA0712400), National Nature Science Foundation of China (NSFC, 61772313 and 11931008), and Interdisciplinary Science Innovation Group Project of Shandong University (2019).

## Author contributions

Q.M. and B.L. conceived the basic idea. Z.L. carried out the computational analysis and data interpretation. L.Z., Q.W., and A.M. designed and drew the figures. D.C. and J.Z. helped polish the manuscript and all the authors wrote the manuscript. All authors read and approved the final manuscript.

## Supplementary Information

### Supplementary information file

This file contains Supplementary Figures S1-3, details in the Problem formulation and Methods part, and Supplementary references.

### Supplementary Tables

This file includes an overview of Supplementary Tables and Supplementary Tables 1-12.

