## Supplemental information for "Inference of disease-associated microbial gene modules based on metagenomic and metatranscriptomic data"

This file includes Supplementary figures S1-3, details in the Problem formulation and Methods part, and Supplementary references.

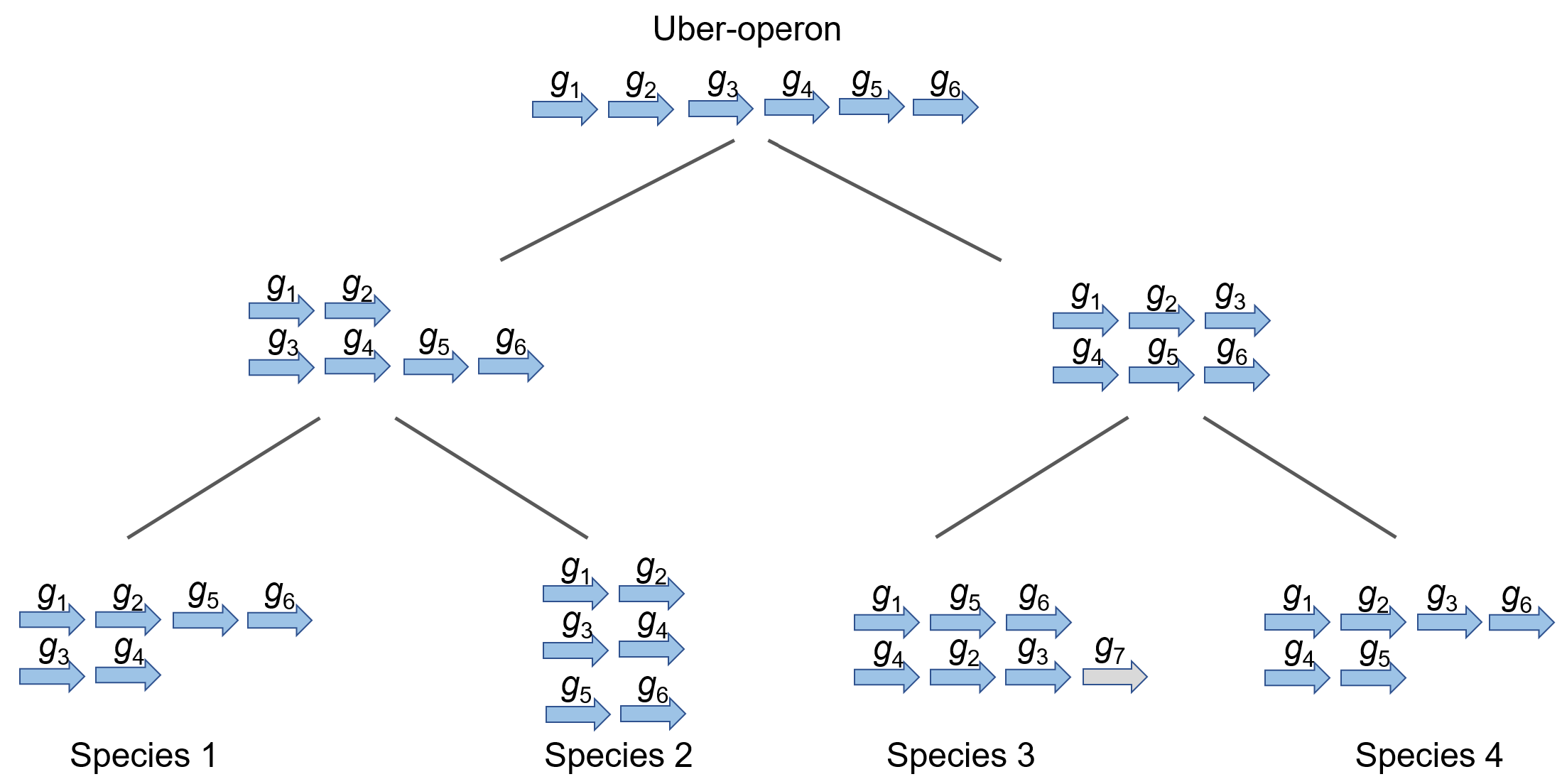

**Fig. S1. A diagram of an uber-operon over evolutionary time.** Each arrow means a gene. The ancestral genome includes six functionally related genes, consisting of an uber-operon. As the population diverges and the genome is rearranged, the functionally related genes are rearranged into new clusters in different species. The grey arrow represents a new gene of related function.

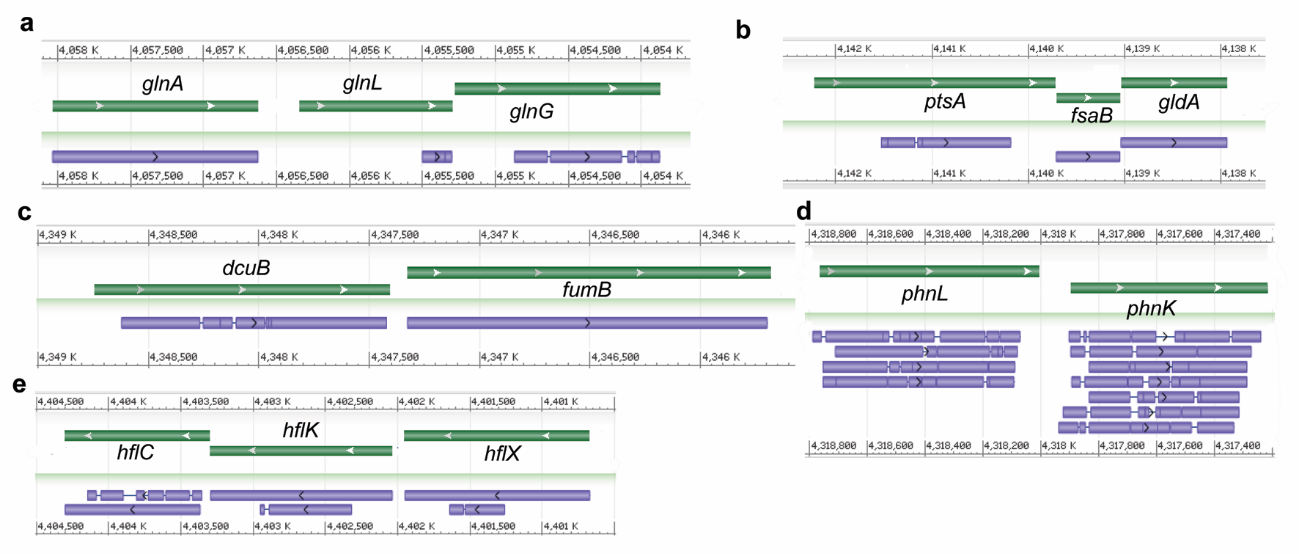

**Fig. S2. The alignment with the representative reference genome of *Escherichia coli*.** **a-e.** Each panel represents an operon from RegulonDB, corresponding to *glnALG*, *ptsA*-*fsaB*-*gldA*, *dcuB*-*fumB*, *phnKL*, and *hflXKC*, respectively. The number text in the head indicates the location of the genome. The green bars with arrows mean the genes on the genome, while the purple bars mean the alignment regions of sequences.

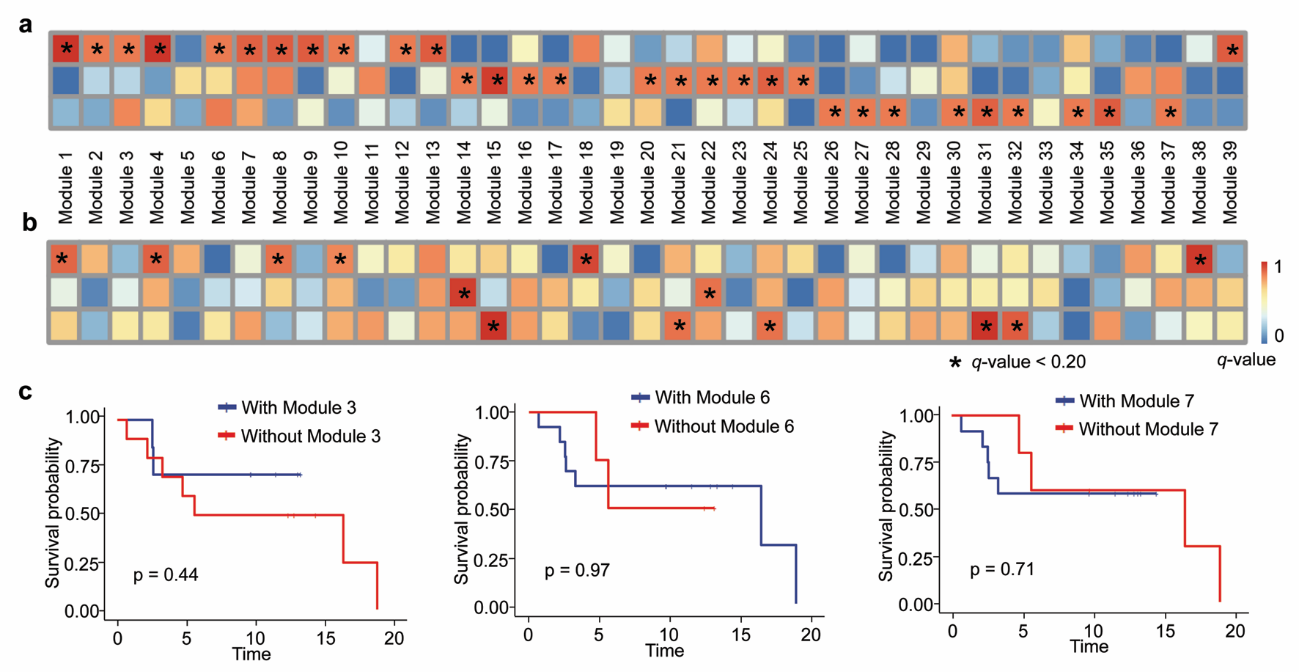

**Fig. S3. The results on the independent data set and the integrated data set. a.** The result of the Wilcoxon signed-rank test on the independent dataset considering modules. **b.** The result of the Wilcoxon signed-rank test considering a single gene. **c.** Kaplan-Meier curves of gene module-associated groups along with corresponding log-rank test *p*-values.

**Section 1. The illustration of the proposed NP-hard problem**

The problem we formulated is to find local low-rank submatrices within expression matrix *A* that maximize the number of connected components between the corresponding gene sets and uber-operons. Simply speaking, the objective is to identify a set of local low-rank submatrices. Construct a weighted graph *G* with genes as nodes, edges linking each pair of genes, and the weight of every edge defined as the number of columns under which the corresponding integers along the two rows are identical. Since the rows will be more similar, the weight of the edge will be heavier. Each of the to-be-identified local low-rank submatrices will correspond to a heavier subgraph on *G* compared to the random ones. But note that not every heavy subgraph corresponds to a local low-rank submatrix. In graph theory, the identification of heavy subgraphs in a weighted graph is NP-hard because the maximum clique problem, a well-known computationally intractable problem, is a special case of this problem (equivalent to the graph with all edges of weight one). Therefore, our problem formulated here is an NP-hard problem.

**Section 2. The construction of the integrated datasets and data pre-processing**

We manually searched publicly available research articles and databases for matched metagenomic and metatranscriptomic data. The PubMed database and the well-curated microbial database, MGnify ^1^, were queried using a collection of relevant keywords, such as “metagenomics”, “metatranscriptomics”, and “disease”. We filtered datasets with missing phenotypes and obtained three datasets associated with melanoma ^2^, type 1 diabetes mellitus/healthy controls without type 1 diabetes mellitus ^3^, and irritable bowel syndrome ^4^, respectively. As a result, 60 samples were obtained and detailed information can be found in Supplementary Table 7. We downloaded the raw sequencing data (.fasta) on NCBI Sequence Read Archive (with a size of about 1.09T). Then, we did data pre-processing of the raw sequencing reads of each sample and constructed the expression matrix. The Kneaddata pipeline (https://huttenhower.sph.harvard.edu/kneaddata) was used for read length filtering, trimming of library primers and adapters, trimming of low-quality read ends, and the removal of host contamination from raw reads. The obtained clean reads of each sample were output to a widely used tool, HUMAnN2 ^5^, to quantify gene expression within the microbial community, during which sequencing depth normalization has been considered. Finally, we incorporated the genes from all samples and relative expression within each sample, forming a gene-sample expression matrix.

**Section 3. The discretization of an expression matrix**

We need to assess the expression similarity of each pair of genes, and the expression value is continuous here. For ease of measuring the similarity of expression for large-scale genes and to reduce the effect of noisy data, we firstly use a qualitative representation and transfer the continuous value into discrete integers ^6^. Specifically, for gene *i*, we sort its expression values within all samples in increasing order:

$$e_{i,1}e_{i,2}\cdots e_{i,s}\cdots e_{i,c-1}e_{i,c}e_{i,c+1}\cdots e_{i,2\times c-s+1}e_{i,2\times c-s+2}\cdots e_{i,n}$$

where *c* = *n*/2, *s* = *n* $\times$ *q* +1, *q* is a parameter set by users. The values that are bigger than $e_{i,c}+d_{i}$ (${d_{i}=min\{e}_{i,c}-e_{i,s}, e_{i,2\times c-s+1}-e_{i,c}$) are thought to be high-expressed, while those smaller than $e_{i,c}-d_{i}$ are low-expressed. Others are of normal expression (represented by the integer 0). Note, since we merely focus on low or high expression, the parameter *q* can be used as 0.5. We further equally partition the high-expressed (sorting in increasing order) and low-expressed values (sorting in decreasing order) into *r* intervals (a parameter which may be pre-specified by the user, generally selection is 1 since this is a rough qualitative representation), then we set all the values belonging to the *j*th interval to be the integer *j and* -*j,* respectively (1$\leq j \leq r$). In this way, the original matrix *A* with continuous values is transferred into a new matrix consisting of integers.

**Section 4. The determination of parameter *w***

The parameter *w* reflects the impact of gene context conservation on function relation between genes. It ranges from 0 to 1. We set multiple different values of *w* (From 0.1 to 0.9, and the step is 0.1) and compared the capacity of identified gene modules to distinguish disease phenotypes (Z-tests in main text Methods). Results can be found in Supplementary Table 12. Based on this, we selected 0.3 as the default value in this article. The validation on the independent dataset and the integrated dataset demonstrated the good performance of IDAM under this parameter.
